# Identification of small-molecule adjuvants that enhance the sensitivity of *Escherichia coli* to nitrofurantoin: Roles of Lon and MarA

**DOI:** 10.64898/2026.01.18.700157

**Authors:** Santhi Sanil Nandini, Pritam Saha, Rithika Shri, Pragyee Sharma, Sirisha Jagdish, Balaram Khamari, Sobhan Sen, Eswarappa Pradeep Bulagonda, Dipankar Nandi

**Author notes:** Corresponding author: FE-14, Department of Biochemistry, Biological Sciences Building, C.V. Raman Road, Indian Institute of Science, Bengaluru-560012, Karnataka, India.

## Abstract

The increasing cases of resistance among UTI pathogens pose a significant threat to the continued clinical use of nitrofurantoin. In this study, we explored the molecular mechanisms underlying nitrofurantoin resistance and investigated the potential of synergistic activity of salicylates in enhancing the antibacterial activity of nitrofurantoin. In our initial observation, deletion of *lon* (Δ*lon*) conferred enhanced susceptibility to nitrofurantoin. We identified the critical role of Lon protease in regulating the sensitivity to nitrofurantoin. Investigation into the mechanisms revealed that the *lon* deletion strains show a higher level of *marA* and *nfsA*, which is likely to facilitate the conversion of nitrofurantoin from its pro form to its active form. The Δ*lon* strains displayed an elevated level of ROS, membrane alteration and filamentation upon treatment with nitrofurantoin. Higher ROS levels and membrane alteration were reversed upon treatment with glutathione, further confirming the role of oxidative stress in mediating the sensitivity to nitrofurantoin. Building on these mechanistic insights, we tested salicylates to synergistically enhance the efficacy of nitrofurantoin by indirectly inducing *marA* through the repression of the mar operon, thereby enhancing nfsA transcription. Both sodium salicylate and acetyl salicylate enhanced the efficacy of nitrofurantoin and lowered the dose of nitrofurantoin required to inhibit the growth of the WT strain. Importantly, this synergistic effect with acetyl salicylate was also observed in nitrofurantoin-resistant clinical isolates, where the combination reduced the effective nitrofurantoin concentration required for growth inhibition. This work provides novel insights into the roles of transcriptional regulators and proteolysis in antibiotic susceptibility, advancing the notion that antibiotic adjuvants are a reliable means of reviving the efficacy of antibiotics.

**Importance:** This study unravels the uncharacterised role of Lon protease in nitrofurantoin susceptibility and illustrates the enhanced efficacy of nitrofurantoin–salicylate combinations as a promising therapeutic strategy to overcome emerging resistance in UTI pathogens. This study highlights the importance of investigating the repurposing of other FDA-approved molecules to combat resistance.

## INTRODUCTION

Antimicrobial resistance is one of the most significant threats faced by the medical world today. The development of resistance in microorganisms towards a wide variety of antibiotics is an alarming situation which needs immediate attention (D’Costa *et al*., 2006). Antimicrobial resistance occurs when microorganisms like bacteria, fungi and viruses develop mechanisms to defeat the drugs designed to kill them (Kumar *et al*., 2013). With the increase in resistance, it becomes difficult to eradicate them, and the drug becomes ineffective. The introduction of new drugs in the market also poses a threat to the emergence of resistance. However, the revival and reintroduction of old antibiotics and their cautious use can cater to this problem to a large extent. The most common multidrug-resistant bacteria in Europe lead to about 400,000 infections and 25,000 deaths annually. Those infections and deaths were attributed to different microorganisms like *S. aureus, E. coli, Enterococcus faecium, Streptococcus pneumoniae, Klebsiella pneumoniae, and Pseudomonas aeruginosa* in 2007 (Prestinaci *et al*., 2015).

Nitrofurantoin is a broad-spectrum bactericidal antibiotic belonging to the class of antibiotics known as “nitrofurans”, and it works against many types of bacteria, including Gram-positive and Gram-negative. It has been widely and successfully used for a long time to treat uncomplicated lower UTIs in adults and children. Nitrofurantoin is effective against many common UTI-causing bacteria, including both Gram-positive species like Enterococcus, Staphylococcus aureus, group B streptococci and Staphylococcus epidermidis, as well as Gram-negative bacteria such as *E. coli, Klebsiella, Shigella, Salmonella, Citrobacter* (Khamari & Bulagonda, 2025). They function by undergoing reduction through nitroreductases. This process also generates intermediates like the nitro anion free radicals and hydroxylamine that non-specifically bind to the ribosome and inhibit critical bacterial enzymes required to synthesise DNA, RNA and protein (Sanni *et al.,* 2024). Following the oral administration, nitrofurantoin is rapidly absorbed in the intestinal tract and subsequently eliminated via renal excretion, leading to high therapeutic concentrations in the urinary system (Mahdizade *et al.,* 2023). Recently, several national and international guidelines, including those from the Infectious Disease Society of America (IDSA), European Society of Clinical Microbiology and Infectious Diseases (ESCMID) and Indian Council of Medical Research (ICMR), have reaffirmed nitrofurantoin as the first-line treatment for uncomplicated lower urinary tract infections (Khamari *et al*., 2022). Nitrofurantoin is widely used, although clinical investigations have shown that it can have adverse side effects, including interstitial pneumonia, liver toxicity, pulmonary fibrosis, and acute pulmonary toxicity (Karpman & Kurzrock, 2004; Goemaere *et al*., 2008; Livanios *et al*., 2016; Sanni *et al*., 2023).

The genes that control nitroreductase activity are known as “nitrofuran sensitivity genes” (*nfsA* and *nfsB*). The *E. coli nfsA* and *nfsB* code for oxygen-insensitive nitroreductases NfsA and NfsB. These enzymes are involved in reducing nitroaromatic compounds and have also been observed to have a bacterial response towards nitrofuran antibiotics. NfsA is a predominant oxygen-insensitive nitroreductase, a flavoprotein tightly bound to flavin mononucleotide (FMN) and has a molecular weight of about 26,799 Da. NfsB is the minor oxygen-insensitive nitroreductase in *E. coli,* which is encoded by *nfsB*, and it can use both NADH and NADPH as electron donors (Zenno *et al.,* 1996). In 1975, a study by the McCalla group found three distinct, separable nitrofuran reductases in *E. coli* and indicated the lack of one such enzyme in some nitrofurazone-resistant mutant strains. Single-step mutants of intermediate nitrofuran resistance possessed the *nfsA⁻ nfsB⁺* genotype and only 20–30% of the wild-type nitroreductases I activity. On the other hand, *nfsA⁺ nfsB⁻* strains still had around 70–80% of the wild-type nitroreductase activity, which was enough to retain wild-type sensitivity to nitrofurans (McCalla *et al*.,1978).

In bacteria, intracellular proteolysis is mediated by ATP-dependent proteases belonging to the vast AAA+ family of proteins, like Lon, FtsH, HslVU, and the Clp family. Lon plays a critical role in the stress response in various bacteria. It enables the pathogens to escape all types of stress, such as heat, oxidative and metabolic stresses. Lon is the most efficient protease in degrading misfolded proteins in cells lacking DnaK and the heat shock proteins in *E. coli* (Chandu & Nandi, 2004). Previous studies from our lab have reported that the deletion of *lon* leads to higher levels of *marA*, as Lon controls MarA levels, and MarA can bind to the mar promoter to upregulate its transcription (Bhaskarla *et al*., 2016; Verma *et al*., 2018; Nandini *et al*., 2025). MarA can bind to the Mar Box upstream of *nfsA* and *nfsB*, thereby controlling their expression (Day *et al*., 2023). In this study, we have explored the roles of Lon and MarA in sensitivity to nitrofurantoin, demonstrating the impact of small molecule adjuvants in reducing the dose of nitrofurantoin and treating nitrofurantoin-resistant pathogens. It is likely that this strategy will reduce the dose of nitrofurantoin.

## RESULTS

### The Δ*lon* strain is resistant to many antibiotics but sensitive to nitrofurantoin compared to the WT

During an MIC broth dilution screening for MIC of different antibiotics for the WT and Δ*lon* strain. We observed that the Δ*lon* strain was resistant to all the antibiotics used in the assay except nitrofurantoin. The MICs of WT were 14 ng/ml, 0.3 μg/ml, 0.75 μg/ml, and 1.4 μg/ml for ciprofloxacin, tetracycline, ampicillin, and nitrofurantoin, respectively. MIC of Δ*lon* was 40ng/ml,0.75 μg/ml,2μg/ml and 0.8μg/ml for ciprofloxacin, tetracycline, ampicillin and nitrofurantoin, respectively. Across different concentrations of the antibiotics, Δ*lon* was resistant to ciprofloxacin and tetracycline (Fig. 1A, SI Fig. 1). To understand the precise nitrofurantoin dose for further experiments, a dose screening was performed where Δ*lon* showed growth reduction in a dose-dependent manner and 0.6μg/ml was selected as the optimal dose for further assays (Fig. 1B). Growth kinetics assay was performed and growth reduction induced by nitrofurantoin was observed till the 6th hr, and growth started to increase post-6th hr slightly, whereas there was a drastic increase in growth post-9th hr (Fig. 1C).This growth reduction is speculated to be an effect of very short half-life of nitrofurantoin or absence of the inactive form of nitrofurantoin in the media post 6 hrs. CFU/mL was also calculated for the 6^th^ hr, and the Δ*lon* strain showed a reduced CFU/mL upon treatment with nitrofurantoin (Fig. 1D).

**Figure 1:**
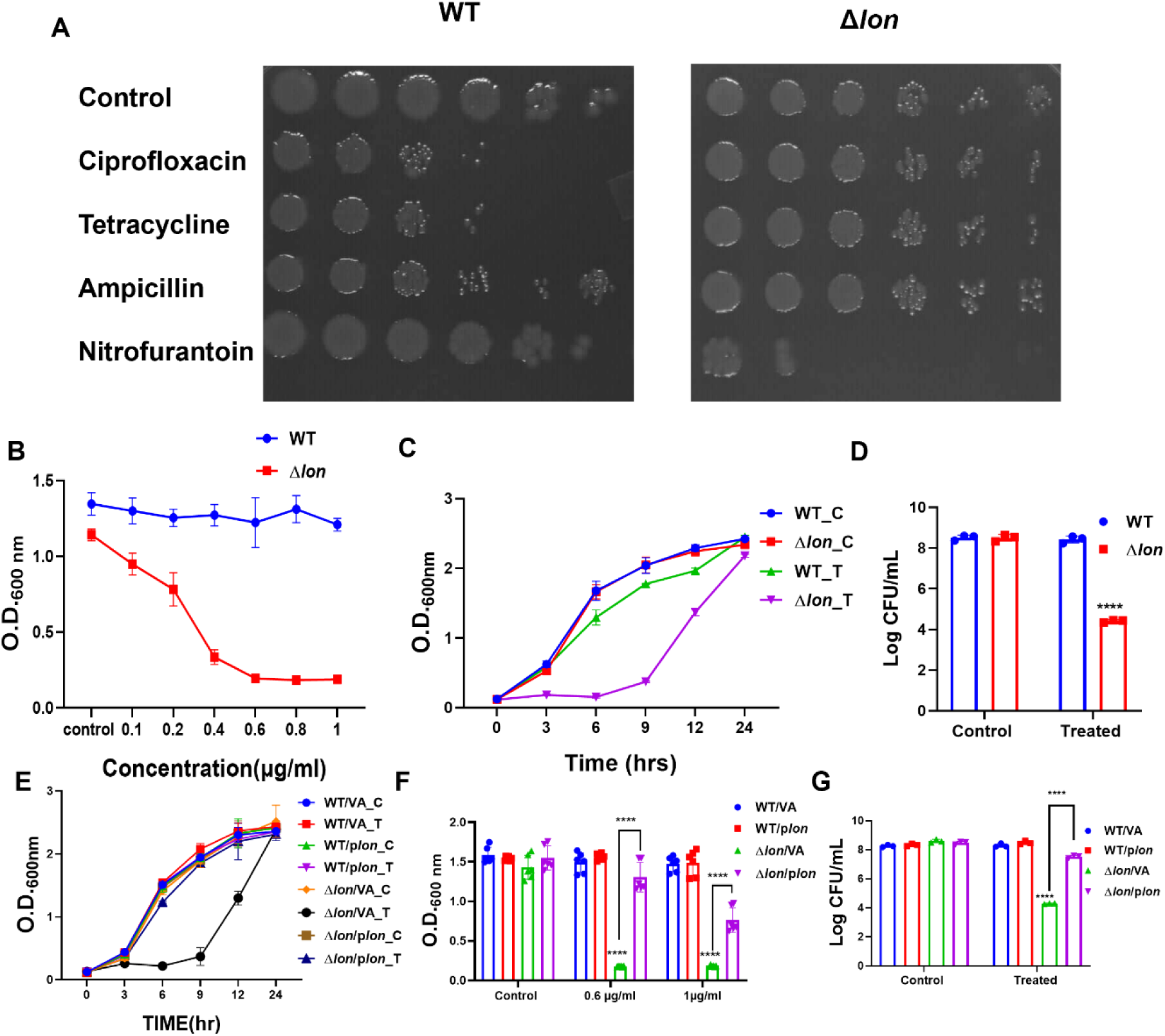
The Δ*lon* strain is resistant to most antibiotics but sensitive to nitrofurantoin and the phenotype is trans-complementable. *E. coli* MG1655 WT and Δ*lon* strains were cultured for 6 hr at 37 °C and 160 rpm in the presence of different antibiotics. (A) Scanned image of agar plates after cultures were serially diluted and replica plated into an agar plate and grown overnight; (B) Growth was assayed by measuring the O.D. at 600 nm using a UV-visible spectrophotometer; (C) Bacterial growth curve plotted as OD/ml v/s time and (D) Bacterial growth at 6^th^ hr plotted as log CFU/ml v/s time showing growth reduction in Δ*lon* strain in the presence of 0.6μg/ml of nitrofurantoin compared to WT; (E) Growth of the strains during exposure to 0.6μg/ml nitrofurantoin; (F) Growth was assayed by measuring the O.D. at 600 nm using a UV-visible spectrophotometer after growing at a low (0.6 μg/ml) and high (1 μg/ml) dose of nitrofurantoin for 6 hr; (G) CFU count is shown 6 hr post-exposure. C represents the control group, and T represents the treated group. The data represent three independent experiments. For statistical analysis, a two-way ANOVA was performed, and the data are plotted as mean ± S.D., where * indicates p < 0.05. Statistical analysis was performed to compare the WT and Δlon strains under different conditions.

### The initial growth reduction in Δ*lon* is rescued by trans-complementation of *lon*

To understand the *lon* dependence of the phenotype observed, *lon* was trans-complemented through the transformation of p*lon*. In the growth kinetics assay, the Δ*lon*/VA strain showed growth reduction till 6^th^ hr, whereas the Δ*lon*/p*lon* strain showed a rescue in the growth reduction, indicating that the growth reduction phenotype observed is *lon* dependent (Fig. 1E). CFU/mL was done at 6^th^ hr, and the rescue in growth upon trans-complementation of *lon* was also observed (Fig.1F). Growth was also assayed with a higher concentration of nitrofurantoin (1μg/ml), showing that the growth reduction in Δ*lon*/VA was rescued in Δ*lon*/p*lon* even at a higher concentration (Fig. 1G).

### Nitrofurantoin induces filamentation of Δ*lon*

To understand the morphological changes occurring in WT and Δ*lon* cells upon NF treatment, atomic force microscopy (AFM) images were recorded for cells treated with nitrofurantoin. Δ*lon* cells showed filamentation, whereas the WT cells did not show any change in cell shape upon treatment with nitrofurantoin (Fig.2A). This is a possible indicator of the SOS response pathway blocking cell division in Δ*lon* cells (Zhang *et al*.,2023).

**Figure 2.**
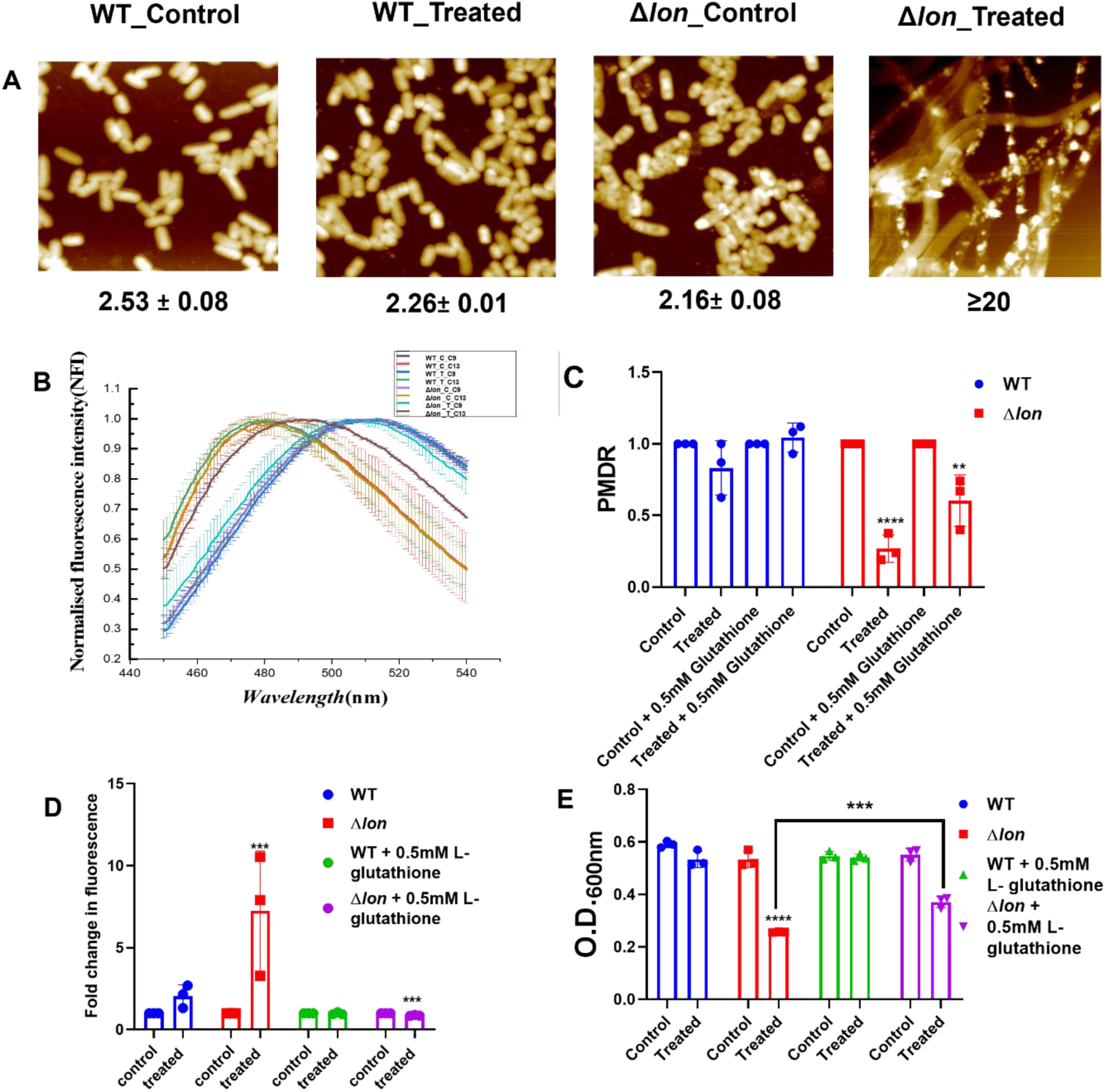
Nitrofurantoin induces filamentation, lowers PMDR and enhances ROS production, which is rescued by glutathione treatment in the Δ*lon* strain. (A) Atomic force microscopic images of WT and Δ*lon* cells grown for 6 hr in the presence of 0.6μg/ml of nitrofurantoin, with cell length mentioned below each image. *E. coli* MG1655 WT and Δ*lon* strains were cultured for 3 hr at 37 °C and 160 rpm in the presence of different concentrations of nitrofurantoin and /or 0.5 mM glutathione. (B) The fluorescence spectra of 4-APC9 and 4-APC13, WT and Δ *lon* cells treated with nitrofurantoin, C represents the control group, and T represents the treated group; (C) PMDR quantified post-treatment with nitrofurantoin and/or glutathione; (D) Fold change in fluorescence plotted after quantifying ROS using DCFDA-based fluorimetric assay; (E) Growth was assayed by measuring the O.D. at 600 nm using a UV-visible spectrophotometer. The data are representative of three independent experiments plotted as mean ± SD. Statistical analysis was performed using two-way ANOVA and one-way ANOVA for (E), where * indicates p<0.05.

### Nitrofurantoin induces membrane alterations and enhances the production of reactive oxygen species

To understand membrane alterations in these strains upon nitrofurantoin treatment, we quantified membrane alterations using an inner membrane binding, solvatochromic 4-aminophthalimide (4-AP) dye 4-APC9 (C9 short alkyl chain) and 4-APC13 (C13 long alkyl chain). These dyes bind at different depths to the inner membrane of bacteria due to their varying tail lengths and hydrophobicity. In a previous study from our lab using these dyes, we demonstrated that upon membrane alteration caused by lipid peroxidation, the peak maxima of 4-APC13 shift towards a lower wavelength. The 4AP-C9, which has a short alkyl chain, exhibits no change in peak maxima during membrane alteration (Dubey *et al*., 2024). As explained in the Materials and Methods section, this change is quantified as the Peak Maxima Difference ratio (PMDR). Upon treatment with nitrofurantoin, the Δ*lon* strain showed a shift in the peak maxima of 4-APC13 to a lower wavelength, indicating membrane alterations. In contrast, the WT strain showed no shift in peak maxima upon treatment with Nitrofurantoin (Fig. 2B). PMDR significantly reduced in the Δ*lon* strain upon nitrofurantoin treatment, but the WT strain did not show any change in PMDR, indicating possible membrane alterations as proposed earlier (Dubey *et al*., 2024). Importantly, treatment with glutathione shows a partial rescue in the PMDR value in the Δ*lon* strain (Fig. 2C).

Conversion of nitrofurantoin to its active form is generally associated with the production of ROS. To understand whether the growth reduction and membrane alterations observed in the Δ*lon* strain result from enhanced ROS production, we quantified ROS production upon nitrofurantoin treatment using a DCFDA-based fluorimetry assay. ROS was quantified at the 3rd hour, as physiological changes are best captured at early time points post-treatment. The Δ*lon* strain showed enhanced fold change in fluorescence of DCFDA upon treatment with NF, whereas the WT did not show any fold change in fluorescence (Fig. 2D). This indicates enhanced ROS production in the Δ*lon* strain and absence of any increase in ROS in the WT strain upon treatment with NF. Treatment with glutathione reduced the ROS levels in the Δ*lon* strain (Fig. 2D). The growth reduction induced by nitrofurantoin was also rescued upon glutathione treatment (Fig. 2E).

### *marA, rob,* and *nfsA* expression is higher upon nitrofurantoin treatment

To gain insights into the mechanism by which nitrofurantoin induces growth reduction in the Δ*lon* strain, we explored the expression profile of key genes using qRT-PCR. The genes coding for nitroreductases (*nfsA* and *nfsB)*, involved in the conversion of nitrofurantoin to its active form, and the genes coding for their transcription factors (marA, rob, and soxS) for the nitroreductase genes (SI Fig. 2) were studied using qRT-PCR. *MarA and Rob* showed a tenfold upregulation in expression. However, *soxS* was not changed upon nitrofurantoin treatment in the Δ*lon* strain (Fig. 3A-C). On the other hand, nitroreductase genes, *nfsA*, showed an eight-fold upregulation in expression, but *nfsB* did not show any change upon treatment with nitrofurantoin in the Δ*lon* strain (Fig. 3D-E). These five genes do not show any change in expression upon treatment in the WT strain. These results indicate that at 0.6μg/ml of nitrofurantoin treatment for 3 hours, increase the expression of *marA, rob* and *nfsA*, but not *soxS* and *nfsB*.

**Figure 3.**
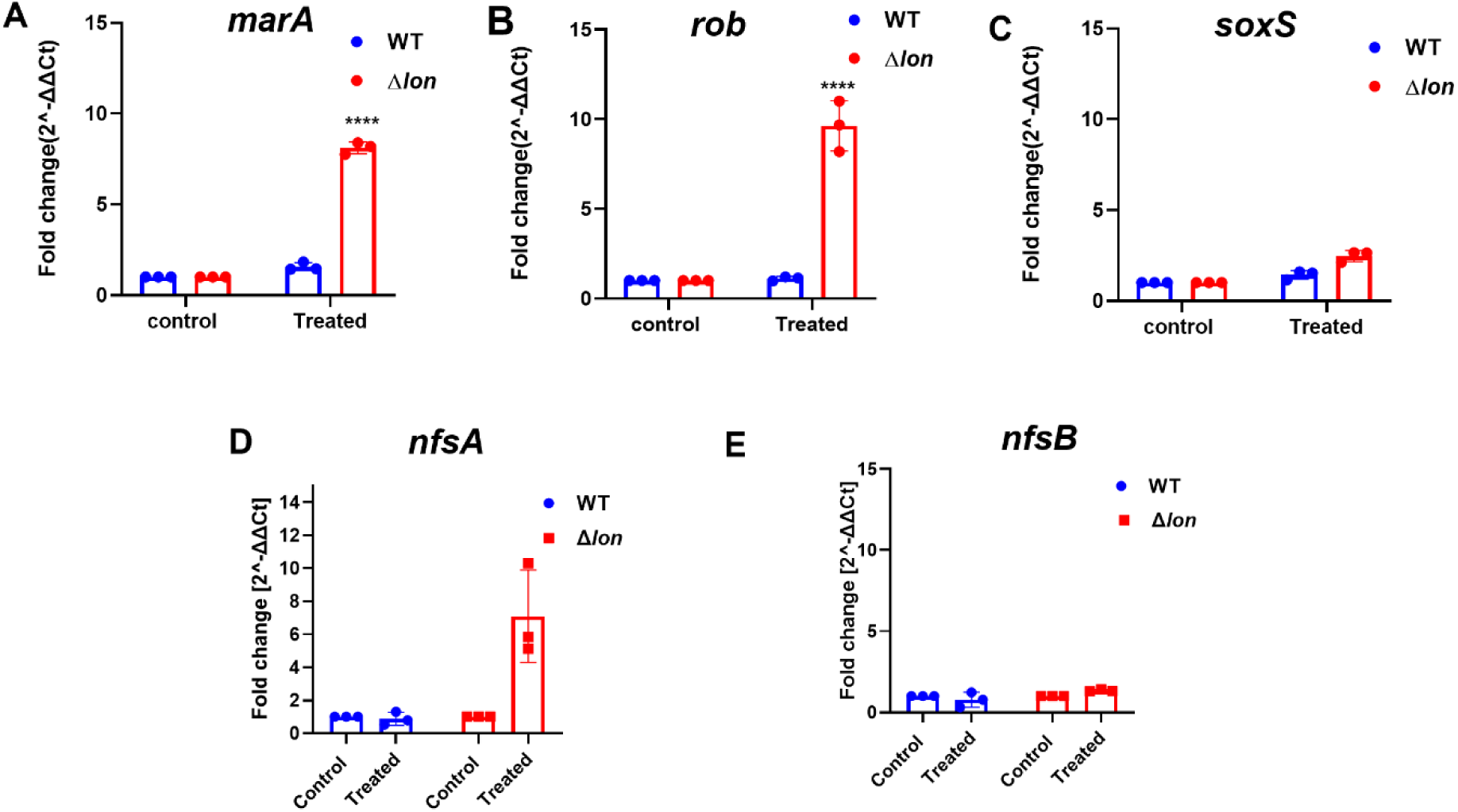
Nitrofurantoin induces expression of *marA, rob*, *nfsA* and *nfsB* but not *soxS.* The cells were grown for 3 hours at 37^○^C, with or without 0.6μg/ml nitrofurantoin; cells were harvested post 3 hr and processed for RNA isolation. Fold changes in transcript levels of (A) *marA*, (B)*rob*, (C) *soxS, (D) nfsA,* and (E)*nfsB* in WT and Δ*lon* strains were determined by quantitative real-time PCR. *gapA* was used as the reference gene. The data is representative of three independent experiments with mean ± SD where * indicates P<0.05. Statistical analysis was performed for each strain relative to its untreated control. Comparison between the strains is indicated wherever significant.

### Overexpression of *marA* lowers the growth of WT strain in the presence of nitrofurantoin

To validate the roles of *marA, rob* and *soxS* in the growth reduction induced by nitrofurantoin in the Δ*lon* strain, knock-out strains of *marA, rob* and *soxS* were screened. The knockouts did not exhibit any growth reduction in the presence of nitrofurantoin, even at a high concentration (SI Fig. 3). This suggests that the absence of these genes does not impact the nitrofurantoin-mediated cell death. Since the growth reduction is speculated to be dependent on the nitroreductases, both genes having a *mar* box upstream, we hypothesised that the overexpression of *marA* in WT would help us unravel the mechanism. Thus, growth kinetics and OD/CFU assays were performed using a *marA* overexpression strain (WT/p*marA*), a *marA* trans-complemented strain (Δ*marA*/p*marA*) and their respective vector plasmid control strains (WT/VA, Δ*marA*/VA) (Fig. 4A). *marA* overexpression strain showed a significant reduction in growth till 6^th^ hr. The growth reduction in WT/p*marA* was also observed in a higher concentration of nitrofurantoin (1μg/ml) (Fig.4 B). Log CFU/ml also showed a reduction in the WT/p*marA* grown in the presence of nitrofurantoin (Fig.4 C). All these results help us to conclude that a higher level of *marA* in the *lon* deletion strain leads to a reduced growth phenotype.

**Figure 4.**
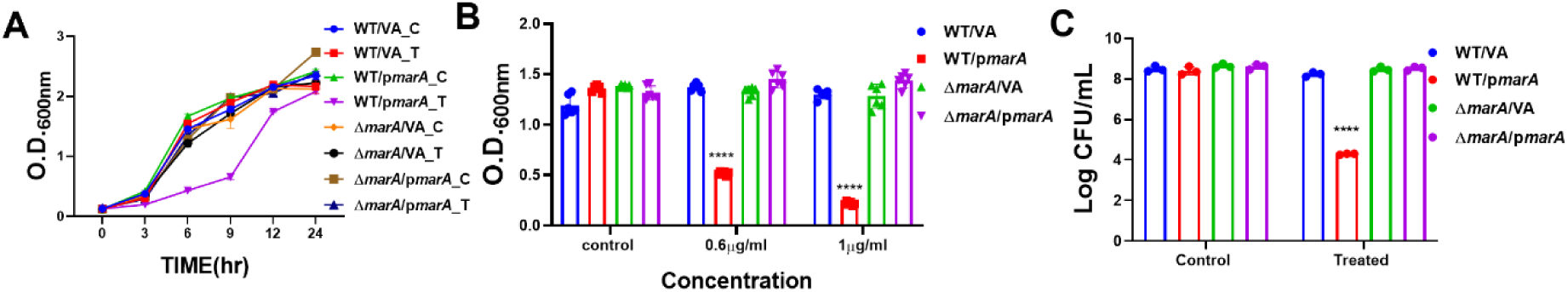
Overexpression of *marA* decreases the growth of WT in the presence of nitrofurantoin. *E. coli* WT /VA, WT /p*marA,* Δ*marA/*VA, and Δ*marA/*p*marA* were cultured in the presence of nitrofurantoin at 370 °C and 160 rpm. (A) Growth of the strains during exposure to 0.6μg/ml nitrofurantoin;(B) Growth was assayed by measuring the O.D. at 600 nm using a UV-visible spectrophotometer after growing the cells at a low (0.6 μg/ml) and high (1 μg/ml) dose of nitrofurantoin for 6 hr; (C) CFU count is shown 6 hr post-exposure; C represents the control group, and T represents the treated group. The data are representative of three independent experiments plotted as mean ± SD. Statistical analysis was performed using two-way ANOVA and one-way ANOVA for (D), where * indicates p<0.05.

### Overexpression of *marA* enhances the sensitivity of Δ*nfsB* strain

Since we have understood the role of *marA* in the growth reduction induced by nitrofurantoin, we wished to explore the role of the nitroreductase genes in the mechanism. A dose screening assay of nitrofurantoin ranging from 0.6μg/ml to 8μg/ml was performed. Growth reduction is observed in the Δ*lon* strain as previously mentioned. WT and Δ*nfsB* strains start showing growth reduction from 6μg/ml, but the Δ*nfsA* strain shows resistance even at a higher concentration of 8μg/ml (Fig. 5 A). Growth kinetics with OD and CFU showed the reduction of growth in the Δ*nfsB* strain till 6^th^ hr (Fig. 5B-C). To further confirm the phenotype, *marA* was overexpressed in Δ*nfsA* and Δ*nfsB*; the vector plasmids were also transformed into these strains (SI Fig 4). Δ*nfsA*/VA, Δ*nfsA*/p*marA* strains did not show any growth reduction in the presence of nitrofurantoin. Δ*nfsB*/VA strain shows growth reduction, which was further lowered in Δ*nfsB*/p*marA,* indicating that the overexpression of *marA* can lead to an increased level of *nfsA* in the strain, leading to growth reduction (Fig.5D). This result also helps us exclude any *nfsB* role in the nitrofurantoin-mediated growth reduction. These results confirm that the absence of the *nfsA* gene in the Δ*nfsA* most likely leads to the failure to convert nitrofurantoin to its active form and thus the resistance phenotype. Overall, the result demonstrates that the NfsA nitroreductase plays a significant role in converting nitroreductase to its active form.

**Figure 5.**
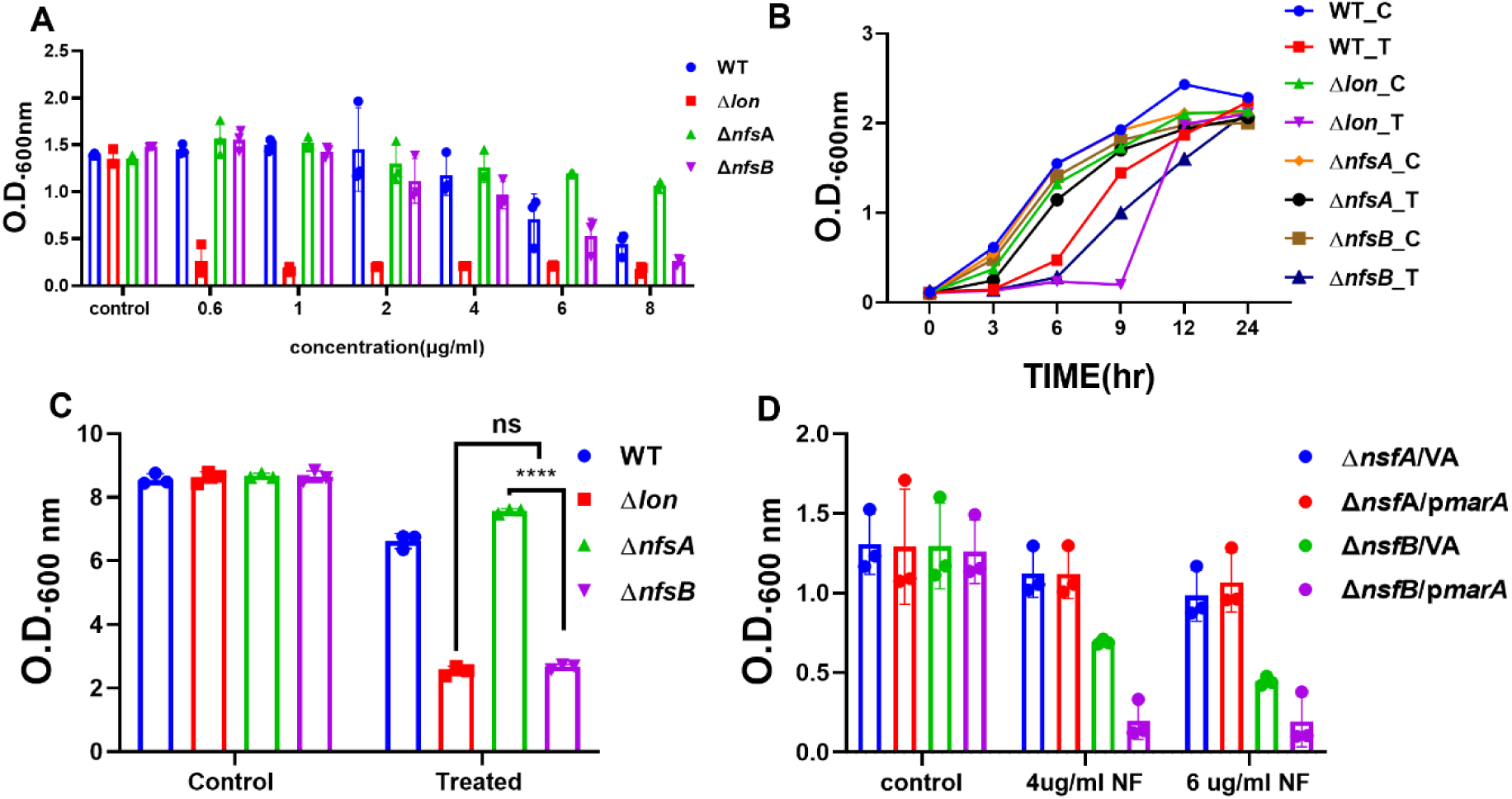
Overexpression of *marA* decreases the growth of the sensitive Δ*nfsB* strain in the presence of nitrofurantoin. *E. coli* WT, Δ*lon,* Δ*nfsA,* Δ*nfsB,* Δ*nfsA*/VA, Δ*nfsB*/VA, Δ*nfsA*/p*marA* and Δ*nfsB*/p*marA* were cultured in the presence of nitrofurantoin at 370 °C and 160 rpm. (A) Growth was assayed by measuring the O.D. at 600 nm using a UV-visible spectrophotometer after growing the cells at different doses of nitrofurantoin for 6 hr; (B) Growth of the strains during exposure to 6μg/ml nitrofurantoin; (C) CFU count is shown 6 hr post-exposure; C represents the control group, and T represents the treated group (D) Growth was assayed by measuring the O.D. at 600 nm using a UV-visible spectrophotometer after growing the cells at different doses of nitrofurantoin for 6 hr. The data are representative of three independent experiments plotted as mean ± SD. Statistical analysis was performed using two-way ANOVA and one-way ANOVA for (D), where * indicates p<0.05.

### Synergistic effect of NF and Sodium Salicylate /Acetyl Salicylate induces growth reduction in WT

The previous results from this study indicate that the overexpression of *marA* can induce growth reduction in WT. Previous studies from our lab have shown that NaSal can induce higher expression of *marA* (Bhaskarla *et al.,*2016). This led us to use NaSal to induce *marA* that may reduce the dose of nitrofurantoin required for inducing growth reduction in WT. We screened the WT and Δ*lon* strains in the presence of nitrofurantoin and/or NaSal. WT in the presence of 4μg/ml of nitrofurantoin or 3mM of NaSal did not show any growth reduction, but in combination with 4μg/ml of nitrofurantoin and 3mM of NaSal, growth was significantly reduced, comparable to the Δ*lon* strain growing in the presence of 4μg/ml of nitrofurantoin alone (SI Fig.5A). Since NaSal is not FDA approved, we used Acetyl Salicylate (AcSal), an FDA-approved salicylate molecule. WT in the presence of 2mM of AcSal did not show any growth reduction. However, the combination of 4μg/ml of nitrofurantoin with 2mM of AcSal showed a significant growth reduction, comparable to the Δ*lon* strain growing in the presence of 4μg/ml of nitrofurantoin alone (SI Fig. 5B). To further confirm the phenotype of cells, flow cytometry was performed, followed by staining of cells with Propidium Iodide (PI) (Rosenberg *et al*., 2019). This helped us to understand the change in morphology and the number of dead cells. Upon treatment with nitrofurantoin, WT showed a 4-fold change in FSC, whereas in combination with AcSal, it increased to 10-fold; SSc showed a 4-fold change with nitrofurantoin alone, and upon combination treatment with AcSal, it increased to 12-fold (Fig. 6). This signifies changes in the cell shape and granularity upon treatment with NF and AcSal. The fold change in fluorescence was approximately 10-fold with nitrofurantoin alone, but upon combination treatment with AcSal, it increased to 60-fold (Fig. 6D). We confirmed this observation using PI, which is taken up by dead cells. The percentage of PI+ cells with nitrofurantoin alone was 10% and it increased to around 60 % upon AcSal combined with nitrofurantoin, which indicated a drastic increase in the number of PI-stained cells.

**Figure 6.**
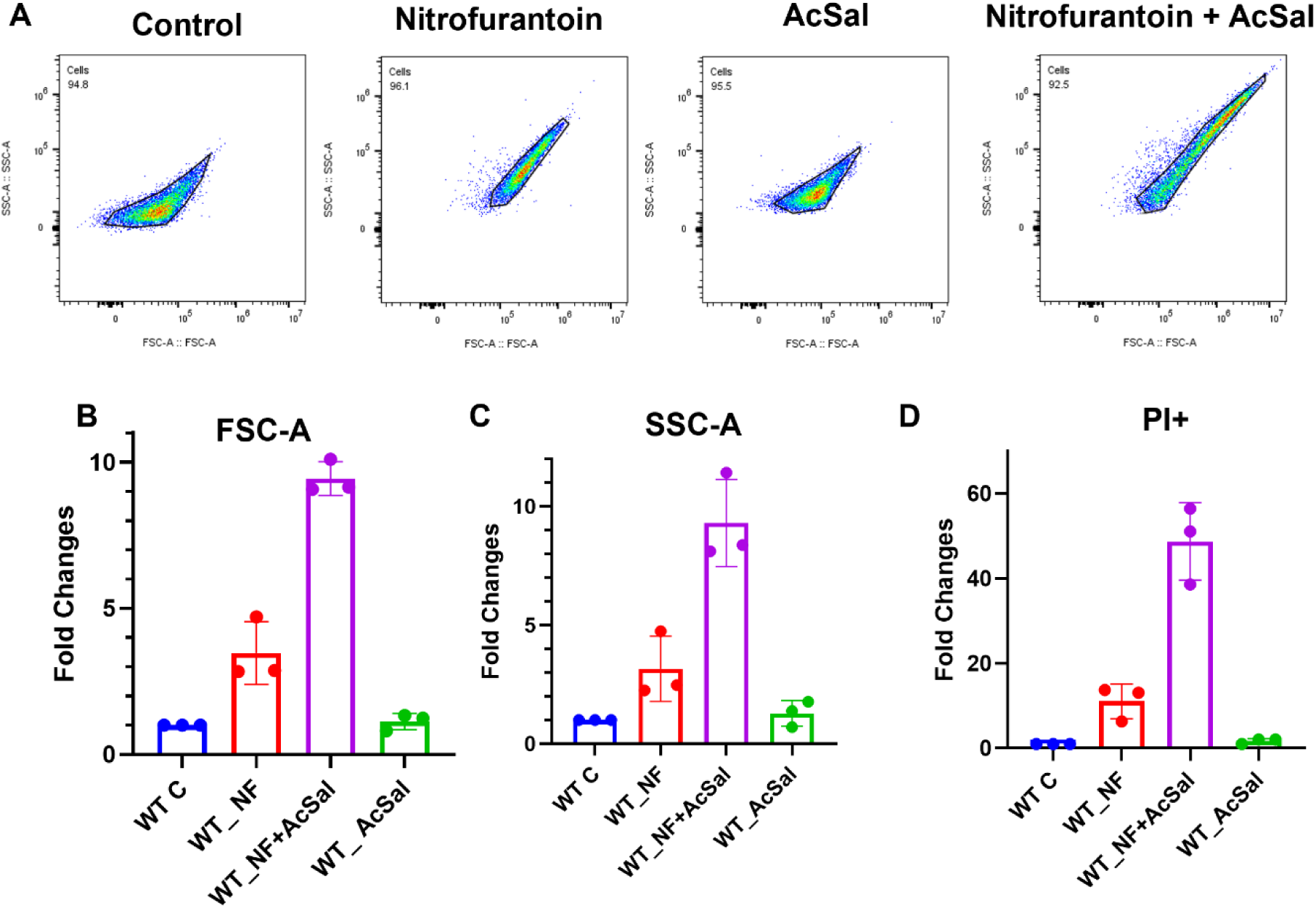
The WT cell population shows a greater shift of FSC-A and SSC-A upon treatment with nitrofurantoin and AcSal. WT cells were cultured in the presence of 4 μg/ml of nitrofurantoin and/or 2 mM AcSal for 3 hours and then washed and stained with PI. (A) Scatter plots showing FSC-A v/s SSC-A under different conditions. Fold change in (C) FSC-A;(D)SSC-A; (E) PI^+^ cells. The data represent three independent experiments, plotted as the mean ± SD. Statistical analysis was performed using two-way ANOVA and one-way ANOVA for (D), where * indicates p<0.05.

### AcSal, in combination with nitrofurantoin, renders sensitivity to clinical strains resistant to nitrofurantoin

It was important to shed light on the above observation in clinical strains of *E. coli*. Two sensitive strains, *E. coli* ATCC 35218 and NS13, and two resistant strains, EC1 and EC3, were used for the assays. An MIC assay was performed to confirm the resistance/sensitivity phenotype of the strains. ATCC 35218 and NS13 showed growth reduction with nitrofurantoin. The EC1 and EC3 strains showed less growth in the control, but the growth didn’t decrease with an increase in the concentration of nitrofurantoin, and this indicates that even though the strains are resistant to nitrofurantoin, they grow slowly compared to the sensitive strains (Fig. 7 A&B).

**Figure 7.**
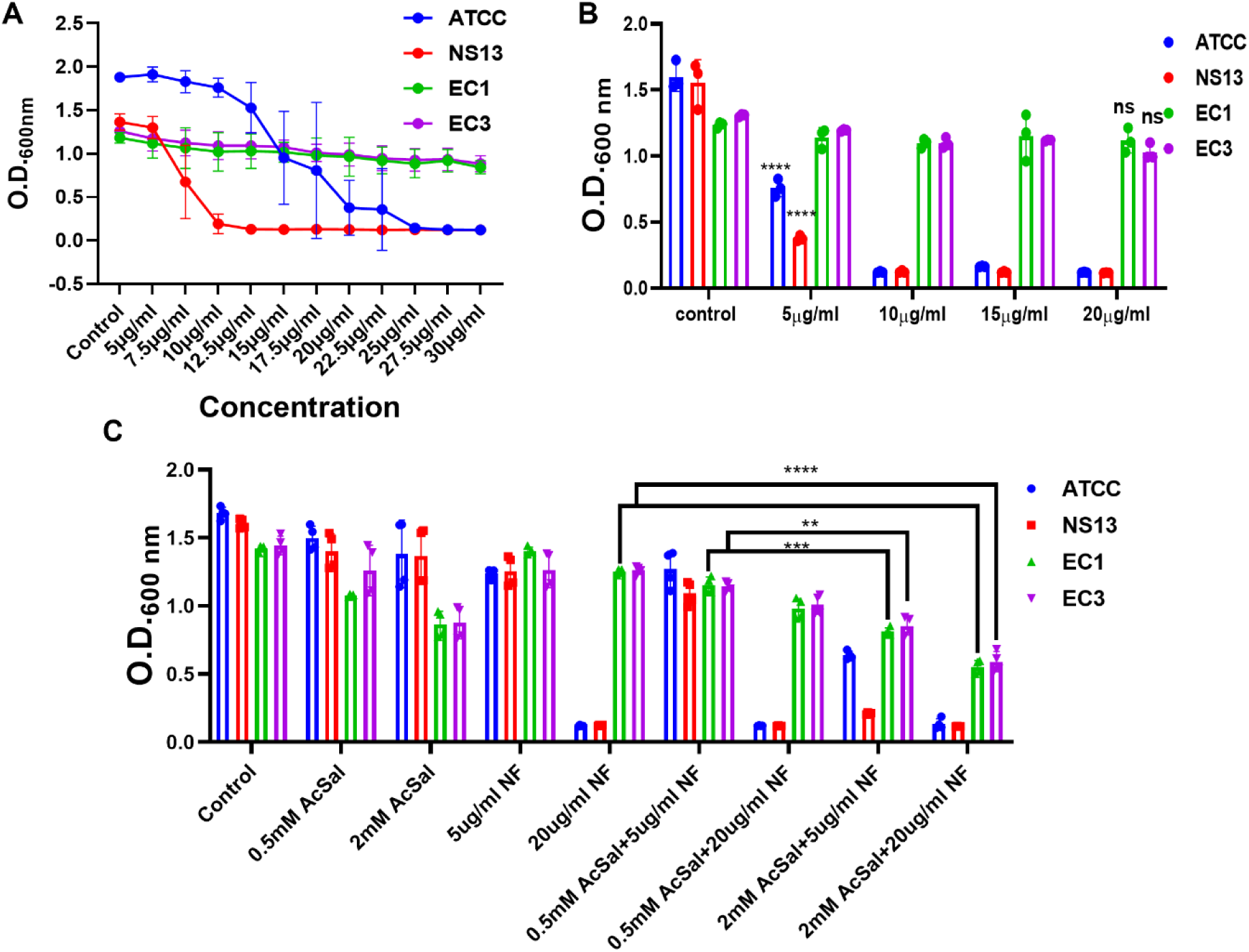
A combination of AcSal with NF induced growth reduction in nitrofurantoin-resistant clinical strains. *E. coli* clinical strains ATCC, EC1, EC3 and NS13 were cultured in the presence of nitrofurantoin at 370 °C and 160 rpm. (A) MIC broth dilution assay with different concentrations of nitrofurantoin; (B) Growth assayed after growing the clinical strains with different doses of Nitrofurantoin; (C) Growth was assayed by measuring the O.D. at 600 nm using a UV-visible spectrophotometer after growing the cells at different doses of nitrofurantoin and AcSal for 6 hr. The data represent three independent experiments, plotted as the mean ± SD. Statistical analysis was performed using two-way ANOVA and one-way ANOVA for (D), where * indicates p<0.05.

To study the synergistic effect of the AcSal-nitrofurantoin combination, the four clinical strains were grown in the presence or absence of nitrofurantoin and/or AcSal. EC1 and EC3 in the presence of 5μg/ml and 20μg/ml of nitrofurantoin or 2mM of AcSal did not show any growth reduction, but in combination of 5μg/ml and 20μg/ml of nitrofurantoin with 2mM of AcSal, growth was significantly reduced, comparable to the sensitive strains ATCC and NS13 growing in the presence of 5μg/ml and 20μg/ml of nitrofurantoin alone, respectively (Fig. 7 C). This indicates the efficacy of the combination in lowering the dose of nitrofurantoin required to reduce the growth of nitrofurantoin-resistant clinical strains.

## DISCUSSIONS AND CONCLUSIONS

This study focuses on the molecular players involved in nitrofurantoin sensitivity and small molecule adjuvants, which aim to enhance the efficacy of nitrofurantoin. The first part of the work explores the role of Lon protease and its substrate MarA and the interplay between *nfsA* and MarA in rendering sensitivity to nitrofurantoin. In the latter part of the work, a combination of salicylates and nitrofurantoin was used to reduce the growth of the WT strain, which was previously resistant to nitrofurantoin, and demonstrated the efficacy of this combination in reducing the growth of nitrofurantoin-resistant clinical strains. In our previous studies, we had shown that Δ*lon* is resistant to most of the antibiotics (Bhaskarala *et al*., 2016; Verma *et al*., 2018; Verma *et al*., 2024). However, in a screening done for this study, we observed a different phenotype with nitrofurantoin; the Δ*lon* strain showed sensitivity to nitrofurantoin, whereas the WT was resistant to it at the concentration used(Fig.1A, SI Fig.1). The MIC and dose screening studies suggested that the sensitivity of the Δ*lon* strain and the sensitivity of the WT strain to nitrofurantoin hold true across different doses (Fig. 3.3 A-D). In the growth kinetics assay, the Δ*lon* strain also showed a slight increase in growth post-6^th^ hr, and this drastically increased after 9^th^ hr (Fig.1C). This increase in growth observed post-6^th^ hr could be due to the short half-life of nitrofurantoin, which is around 20-30 minutes (Mahdizade *et al*., 2023). It can also be due to the absence of nitrofurantoin, which can be converted to the active form along with the release of ROS, since this step plays a key role in nitrofurantoin-mediated killing of bacterial cells (Wang *et al*., 2008; Kaye *et al*., 2025). We also observed that this phenotype can be complemented through trans complementation of *lon*, confirming the complete *lon* dependence of the growth reduction phenotype observed in the Δ*lon* strain (Fig 1E-G).

Morphological changes are commonly observed in bacterial cells as part of coping with stress (Shen *et al*., 2016; Cushnie *et al*., 2016; Verma *et al*., 2018). To understand the morphological changes, atomic force microscopy was employed (Li *et al*., 2010). We observed a nitrofurantoin-mediated cell filamentation in the Δ*lon* strain but not in the WT, which is a novel finding with nitrofurantoin (Fig.2A). A similar filamentation was previously reported with ciprofloxacin in many studies (Bos *et al*., 2015; Bhaskarala *et al*., 2016; Verma *et al*., 2018). Filamentation of cells mediated by ciprofloxacin occurs when the antibiotic interferes with the normal cell division (Galina *et al*., 2017). Ciprofloxacin inhibits DNA gyrase, leading to the switching on of the SOS response, a DNA repair mechanism contributing to filamentation. ROS production during the reduction of nitrofurantoin is a known phenomenon, and this could be leading to DNA damage, SOS response and thus filamentation of cells (Qin *et al*.,2015). In a previous study, collateral sensitivity to nitrofurantoin was observed in spontaneous mutants with a mutation in *lon.* The study also claims that switching on the SOS response and SulA abundance play a major role in the sensitive phenotype observed (Roemhild *et al*., 2020). To understand if nitrofurantoin induced membrane damage, we explored nitrofurantoin-mediated membrane perturbations using solvatochromic interfacial 4-aminophthalimide dyes (4-AP). In our previous study, we used these dyes to differentiate between sensitive and resistant strains of different Gram-negative bacteria, since most of the antibiotics are known to induce ROS and contribute to lipid peroxidation of the membrane, leading to membrane damage (Dubey *et al*., 2024; Singh *et al*., 2016). Two 4-AP dyes, 4AP-C9 and 4-APC13, were used for the study, and these dyes shed light on membrane damage. We have observed that nitrofurantoin-mediated membrane damage in the Δ*lon* strain, further rescued upon glutathione treatment (Fig. 2B & C). These observations shed light on different mechanisms by which nitrofurantoin renders sensitivity to the Δ*lon* strain. To understand if there is an elevated level of ROS in Δ*lon*, a DCFDA-based fluorimetry was performed (Fig. 2D). We observed elevated levels of ROS in the *lon* deletion strain, which were reduced upon treatment with glutathione (Fig. 2D). This result confirms filamentation and membrane damage induced by nitrofurantoin in the *lon* deletion strain could be due to elevated ROS.

Next, we wanted to explore the key genes involved in the growth reduction. qPCR aided us in understanding the key genes involved in the mechanism. Nitrofurantoin is known to be reduced to its active form by nitroreductases encoded by two nitroreductase genes, *nfsA* and *nfsB* (McOsker *et* al., 1994; Day *et al*., 2023). Both these genes have a *mar* box upstream of them, which controls their transcription. Transcription factors MarA, Rob and SoxS are known to bind to these types of *mar* box, regulating transcription of these genes (Chubiz, 2023) (SI Fig. 2). These three are transcription factors whose levels in the cell are controlled by Lon protease (Kirthika *et al*., 2023). We speculated that the absence of Lon protease in the *lon* deletion strain could lead to disrupted homeostasis of these proteins, leading to increased sensitivity to Nitrofurantoin. In qPCR results, we observed positive upregulation of *marA*, *rob* and *nfsA* but not *soxS* or *nfsB* in the *lon* deletion strain. These results unveil the role of the upregulated genes in the sensitivity to nitrofurantoin (Fig. 3). To further confirm the phenotype, we screened strains lacking *marA, rob* and *soxS* but failed to find any role in sensitivity to nitrofurantoin (SI Fig. 3). Previous studies have also reported that knock out of *marA* doesn’t affect the growth in presence of nitrofurantoin (Roemhild *et al*., 2020), but the role of overexpression of *marA* associated with growth reduction observed in Δ*lon* remained unexplored. We have addressed this caveat and established the role of *marA* overexpression in growth reduction by nitrofurantoin. WT/p*marA* strain indeed showed growth reduction in the presence of nitrofurantoin, implying the role of increased level of MarA in growth reduction induced by nitrofurantoin (Fig. 4). Since the upregulation of *nfsA* is correlated with the upregulation of *marA*, a *marA* overexpression system was used to study if overexpression would exhibit any specific phenotype (Fig.5D). These findings suggest that the reduced growth of the *lon*-deficient strain is due to an increased amount of MarA.

Previous studies have shown that nitrofurantoin can cause adverse effects such as peripheral neuropathy in patients with impaired renal function when prescribed for extended periods (Hanlon *et al*.,2019). The use is also restricted in patients with conditions such as bacterial pyelonephritis, anuria, oliguria, or significant renal impairment. Additionally, it is not prescribed for treating UTIs in pregnant women (38–46 weeks of gestation), neonates, men with prostatitis, and elderly patients (over 65 years old) due to the risk of hemolytic anemia, nitrofurantoin’s limited ability to penetrate prostatic tissue effectively, as well as potential pulmonary toxicity, hepatotoxicity, and peripheral neuropathy (Squadrito, 2023). While nitrofurantoin resistance has historically been low, there is a trend of steady increase in resistance over the past few decades (Osei, 2018). In one of the previous studies from our lab, we have shown that NaSal can induce *marA* (Bhaskarla *et al*., 2016). With this knowledge, we decided to induce *marA* with NaSal to check if it can help reduce the dose of nitrofurantoin required to induce growth reduction in WT. 3mM NaSal in combination with 4μg/ml of nitrofurantoin was able to reduce the growth of WT, even though nitrofurantoin alone at this concentration was not able to induce growth reduction (SI Fig.5A). NaSal is not FDA-approved as a standalone drug according to the US Food and Drug Administration, so we decided to check if another salicylate can induce the same effect in combination with nitrofurantoin. AcSal was used at 2mM concentration and was effective in reducing growth in combination with nitrofurantoin at 4μg/ml (SI Fig. 5B). The AcSal mediated cell death and morphological change due to ROS production was further confirmed using flow cytometry. FSC-A and SSC-A showed greater fold change with nitrofurantoin and AcSal combination (Fig. 3. 14). PI staining percentage increased from 10% in nitrofurantoin treatment to 60% in nitrofurantoin and AcSal combination treatment (Fig. 6).

As we have proved the efficacy of this combination in reducing the concentration of nitrofurantoin required to reduce the growth of the WT strain of *E. coli*, we decided to test the efficacy of this combination in clinical strains. Four different clinical strains, two of which are sensitive to nitrofurantoin and two of which are resistant to nitrofurantoin, were screened against the combination (Fig. 7 A-C). This assay revealed the efficacy of the combination in reducing the dose of nitrofurantoin needed to reduce the growth of nitrofurantoin-resistant clinical strains. This also calls attention to the importance of studying FDA-approved molecules like Acetyl salicylate for repurposing.

In conclusion, our study explores the mechanistic framework involving the Lon, MarA and NfsA interaction as well as a therapeutic approach for enhancing the efficacy of nitrofurantoin by employing small molecule adjuvants. Our results demonstrate that the loss of Lon results in an enhanced level of MarA, further leading to an increased level of NfsA, converting more of the pro form of nitrofurantoin to the reduced active form. This, in turn, increases intracellular ROS, causing membrane perturbations, induction of SOS responses and subsequent filamentation of *lon* deletion strains. These findings highlight the role of oxidative stress in mediating sensitivity to nitrofurantoin. By exploring regulatory interactions between Lon protease, transcriptional activator (MarA), and nitroreductase gene (*nfsA*), we reveal novel molecular determinants of nitrofurantoin susceptibility that have not been previously characterised (Fig 8).

**Fig 8.**
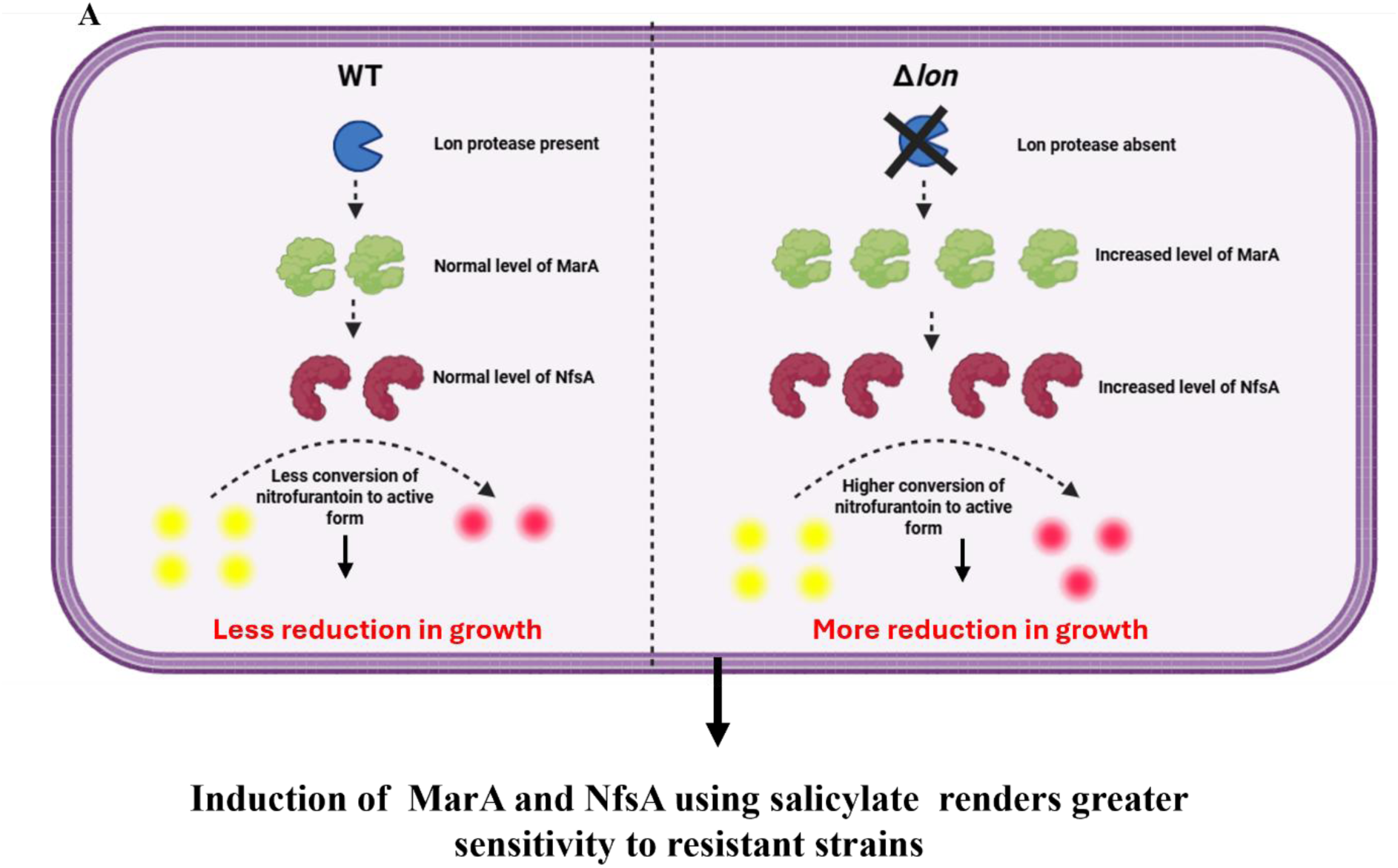
A schematic model representing the roles of Lon, MarA and NfsA in inducing sensitivity to nitrofurantoin. (A). The roles of Lon protease and MarA in inducing NfsA and rendering sensitivity to nitrofurantoin are shown (Created in Biorender).

Furthermore, our findings suggest that the Nitrofurantoin-salicylate combination, through MarA-induction, can significantly reduce the dose of nitrofurantoin in the *E. coli* WT strain. Importantly, this synergistic effect was validated in nitrofurantoin-resistant clinical strains, demonstrating the translational potential of this approach. By lowering the effective dose through an adjuvant-based approach, the adverse effects associated with high doses of nitrofurantoin could be potentially reduced. This approach also helps to overcome the resistance associated inefficiency of nitrofurantoin. From a translational perspective, this work highlights a therapeutic strategy to expand the clinical use of nitrofurantoin, particularly considering its rising resistance rates and the side effects associated with its long-term use. Overall, our research not only offers new mechanistic insights into the interactions among transcriptional regulation, antibiotic susceptibility, and bacterial proteolysis, but it also advances the idea that antibiotic–adjuvant combinations are a practical way to rejuvenate the effectiveness of antibiotics against pathogens that have developed resistance.

## Materials and Methods

### Bacterial strains and growth conditions

The bacterial strains used in this study are listed in Table S1. All the cultures were grown in Luria Broth (LB) comprising Tryptone (10 g/L; HiMedia, Mumbai, India), NaCl (10 g/L; HiMedia) and yeast extract (5 g/L; HiMedia) at 37°C and 160 rpm. Overnight-grown cultures obtained from a single colony of the strains served as the inoculum for all experiments. Antibiotics were used at 100 μg/ml ampicillin, 30 μg/ml chloramphenicol, and 50 μg/ml kanamycin (HiMedia). The optical density (O.D.) of all the strains was normalised to O.D. 2, corresponding to 10^9^ CFU/ml, before all experiments, and 0.2% of these O.D. 2 cultures were used as the starting inoculum for all experiments. A UV-visible spectrophotometer (Tecan, Männedorf, Switzerland) determined growth by measuring the O.D. at _600 nm_.

### Spotting assay

Bacterial cultures were serially diluted and spotted on solid LB agar after growing in the presence or absence of 20ng/ml ciprofloxacin, 1.5μg/ml tetracycline,2μg/ml ampicillin or 0.6μg/ml of nitrofurantoin for 6 hr at 37°C and 160 rpm in the dark. Growth was monitored visually on plates post 15 hr of incubation at 37°C (Bhaskarala *et al*.,2016).

### Determination of Minimum Inhibitory Concentration (MIC)

MIC for the bacterial strains was performed using the broth dilution method. The bacterial cultures grown overnight at 37⁰C and 160rpm served as the pre-inoculum. 200μl of the LB was aliquoted in a 96-well microtiter plate (Tarsons, Kolkata, India), followed by the addition of different concentrations of ciprofloxacin (HiMedia), ranging from 5 ng/ml to 50 ng/ml and tetracycline (HiMedia), ranging from 0.1μg/ml to 2.5 μg/ml, ampicillin (HiMedia), ranging from 0.1μg/ml to 2.5μg/ml and Nitrofurantoin (Sigma), ranging from 0.1μg/ml to 1.8μg/ml. All the wells used 0.2% of OD2 culture as an inoculum. The plate was then incubated at 37℃ for 18 hours, followed by measurement of O.D. values at 600nm. The antibiotic concentration that reduced growth to half was recorded as the MIC value (Nandini *et al*., 2025).

### Atomic force microscopy (AFM) imaging

Bacterial cultures were grown for 6 h at 37°C, under different conditions, after which they were centrifuged at 5000 Xg for 5 min. The cell pellets were washed three times with sterile MilliQ water to remove residual media components. The OD was set to 0.2, and 100μ μL of cell suspension was added to a glass coverslip. The samples were then air-dried at room temperature in a laminar flow hood overnight. The non-adherent bacteria were removed by washing the coverslip twice with MilliQ water, followed by air-drying for 1 hr. The coverslips were then fixed onto a magnetic stub using double-sided carbon tape and transferred to the AFM stage for imaging. All AFM measurements were performed using an NX-10 atomic force microscope (Park Systems). A silicon AFM probe tip (Acta; Park Systems) was used with a resonating frequency of 300kHz.The AFM instrument was operated in the non-contact mode in a scan area 20μm^2^. A pyramidal cantilever 125 mm in length, 35 mm in width, and 4.5 mm in thickness, with a tip radius of 10 mm, was used (Verma *et al*., 2018).

### Flow cytometry

Bacterial cultures were grown for 3 hrs in the presence or absence of 4μg/ml of nitrofurantoin and/or 2mMAcSal in the presence or absence of 0.6μg/mL nitrofurantoin, normalised to O.D of 0.5 and centrifuged at 7000 rpm for 5 minutes, followed by two washes in 1X phosphate-buffered saline (PBS) and stained with 1μg/ml of Propidium Iodide (PI) for 5 mins. A CytoFLEX LX machine was used to record forward scatter (FSC), side scatter (SSC) and cell fluorescence with a Y5B5-PE-A bandpass filter, with 10,000 events analysed per sample. Analysis of the obtained data was performed using the FlowJo software (Dubey *et al*.,2024).

### Cloning of *marA* for trans-complementation and overexpression

The calcium chloride method was used to prepare competent cells. Bacterial cultures were grown overnight from glycerol stock in the presence of antibiotics specific to the resistance cassette for the pre-inoculum. The pre-inoculum was normalised to O.D. 2, and 0.2% of O.D. 2 cells were cultured in 50ml flasks till the mid-log phase (O.D. 0.4 – O.D. 0.5). The flask was incubated at 4°C for one and a half hours, 15 mL of the cell culture for 15 mins at 4°C. The cells were resuspended in 7.5ml of filter-sterilised 100mM MgCl_2_ (HiMedia), incubated at 4°C for 20 mins, and then pelleted down at 3000xg for 15 mins at 4°C. Following this, the cells were resuspended in 7.5ml of filter-sterilised 100mM CaCl2 (HiMedia), incubated at 4°C for 20 mins, and then pelleted down at 3000 Xg for 15 mins at 4°C. Then, the cells were gently resuspended in 1 ml of 100 mM CaCl_2_ in 15% glycerol.100μL aliquots were transferred to Eppendorf, flash-frozen with liquid nitrogen and stored at –80°C.

The WT MG1655 was used as the template for amplifying the *marA* gene using specific primers (Table S2) and Taq DNA polymerase (G-biosciences, Delhi, India). *marA* was targeted for cloning between EcoRI and HindIII sites in the pQE60 plasmid using T4 DNA ligase (DX/DT, Bengaluru, India). The ligated pQE60 plasmid with the *marA* gene was amplified using the positive clones. The heat-shock method transformed the positive clones and control vectors into WT, ΔmarA, Δ*nfsA* and Δ*nfsB* (Nandini *et al*.,2025).

### Quantification of ROS

For the quantification of Reactive Oxygen Species (ROS), bacterial cultures were grown for 3 hr in the presence or absence of 0.6μg/mL nitrofurantoin, normalised to O.D of 0.5 and centrifuged at 7000 rpm for 5 minutes, followed by two washes in 1X phosphate-buffered saline (PBS) and stained with 2μM 2′,7′-Dichlorodihydrofluorescein diacetate (DCFDA) for 30 minutes at room temperature in the dark. The stained cells were washed with PBS, and 200μl of each sample was transferred to a black 96-well plate. The amount of ROS was measured using fluorimetry, with an excitation wavelength of 495 nm and an emission wavelength of 525 nm (Dubey *et al*.,2024).

### Quantification of membrane alterations using 4-APC9 and 4-APC13 dyes

For quantifying membrane alterations, bacterial cultures were grown for 3 hr in the presence or absence of 0.6μg/mL nitrofurantoin, normalised to an O.D. of 0.5 and centrifuged at 7000 rpm for 5 minutes, followed by two washes in 1X PBS before being fixed with paraformaldehyde (PFA) at 37⁰C for 30 minutes. After fixation, cells were washed with 1X PBS 2 times and stained with 50μM of C9 and C13 in the dark for 30 minutes. After a final PBS wash, 200μl of each sample was transferred to a black 96-well plate. Membrane alterations were then assessed by scanning fluorescence spectrum with an excitation wavelength of 375nm and an emission wavelength ranging from 450nm to 540nm (Dubey *et al.,* 2024). The membrane alterations were quantified by calculating the Peak Maxima Difference Ratio (PMDR) by comparing the peaks of AP-C9 and AP-C13 dyes under antibiotic treatment using the formula **PMDR = (Peak** _C13_**-Peak** _C9_) _Treated_ **/ (Peak** _C13_**-Peak** _C9_) _Control_ (Treated =treated with nitrofurantoin (Dubey *et al*.,2024).

### RNA purification, cDNA synthesis, and qRT-PCR

Total RNA was extracted from the WT and Δ*lon* cells grown for 3 hr in the presence or absence of 0.6μg/ml of nitrofurantoin at 37°C and 160 rpm in the dark. Briefly, the cells were lysed and RNA was extracted using the isoPwr-STM total RNA isolation kit (NGIVD, Haryana, India). RNA concentration and purity were quantified using a NanoDrop Spectrophotometer (Thermo Scientific, Waltham, MA, USA), followed by DNase treatment. Subsequently, the RNA was reverse transcribed to cDNA using Revert Aid (Thermo Scientific, Waltham, MA, USA). DNA contamination was tested by PCR amplification of *rrsC*. qPCR was carried out using the Bio-Rad CFX Connect System (Bio-Rad, CA, USA). The primers used are listed in Table S3. GapA was used as the reference gene for all conditions, and fold change was calculated using the 2^−ΔΔCt^ method. The primer efficiency for each primer pair was determined before performing PCRs, according to the MIQE guidelines (Livak & Schnittger, 2001).

### Statistical analysis

All statistical analyses were performed using ANOVA and GraphPad Prism 8, wherever applicable. Data were represented as mean ± S.D., where * denotes p ≤ 0.05, ** denotes p ≤ 0.01, *** denotes p ≤ 0.001, and **** denotes p ≤ 0.0001.

