## Supplemental Information for "Identification of small-molecule adjuvants that enhance the sensitivity of *Escherichia coli* to nitrofurantoin: Roles of Lon and MarA"

**SI Table 1: List of strains used in the study**

| Sl.No | Strain | Genotype/feature | Reference |
|---|---|---|---|
| 1 | WT MG1655 | λ, rph-1 | Guyer *et al*. (1981) |
| 2 | Δ*lon* | MG1655, *lon* : : *cat* | Bhaskarla *et al*. (2016) |
| 3 | Δ*marA* | MG1655, *marA* : : *kan* | Baba *et al*. (2006) |
| 4 | Δ*lon*Δ*marA* | MG1655, *lon* : : *cat*, *marA* : : *kan* | Bhaskarla et al. (2016) |
| 5 | Δ*acrB* | MG1655, *acrB* : : *kan* | Baba *et al*. (2006) |
| 6 | WT/VA | Isogenic complement strain for WT with plasmid pQE60, $Amp^r$ | This study |
| 7 | WT/p*marA* | Isogenic complement strain for WT expressing marA under T5 promoter present in plasmid pQE60, $Amp^r$ | This study |
| 8 | Δ*marA*/VA | Isogenic complement strain for Δ*marA* with plasmid pQE60, $Amp^r$ | This study |
| 9 | Δ*marA*/p*marA* | Isogenic complement strain for Δ*marA* expressing marA under T5 promoter present in plasmid pQE60, $Amp^r$ | This study |
| 10 | *E.coli* Δ*lon*::*kan* | *lon*-deficient derivative of wild type | Kind gift from Dr. Nishad Matange, IISER (Pune, India) |
| 11 | *E.coli* Δ*nfsA* | *nfsA*-deficient derivative of wild type | Kind gift from Prof. Amit Singh, IISc (Bengaluru, India) |
| 12 | *E.coli* Δ*nfsB* | *nfsB*-deficient derivative of wild type | Kind gift from Prof. Amit Singh, |

| | | | |
|---|---|---|---|
| | | | IISc (Bengaluru, India) |

**SI Table 2: List of plasmids used in the study**

| Sl No | Plasmid | Description | Reference |
|---|---|---|---|
| 1 | pQE60 | Low copy bacterial expression plasmid used for trans complementation from the constitutive T5 promoter. It contains an Ampicillin resistance cassette | Chandra *et al.*,2017;2020 |
| 2 | pBAD33 | Low copy number expression vector regulated by the arabinose operon | Matange,2020 |
| 3 | pBAD33-Lon | Plasmid for expression of Lon protease from an arabinose inducible promoter (Matange,2020) | Matange,2020 |

**SI Table 3: List of primers used in the study**

| **Sl.No** | **Primer name** | **Sequence (5'-3')** |
|---|---|---|
| 1 | qRT *gapA* FP | TTTCCGTGCTGCTCAGAAAC |
| 2 | qRT *gapA* RP | GTCAACACCAACTTCGTCCC |
| 3 | qRT *nfsA* FP | GAACTTATTTGTGGCCATCG |
| 4 | qRT *nfsA* RP | TCACCAGTTCTTCACGTAAC |
| 5 | qRT *nfsB* FP | GGTTTATCTCAACGTCGGTA |
| 6 | qRT *nfsB* RP | GGTGTAGCCTTTCTCTTTCA |
| 7 | qRT*marA* FP | TGTCCAGGACGCAATACTGACG |

| | | |
|---|---|---|
| 8 | qRT *marA* RP | TTTTGAAGGTTCGGGTCAGA |
| 9 | Comp *marA* FP | CCGGAATTCATGTCCAGACGCAATACTGACG |
| 10 | Comp *marA* RP | CCCAAGCTTGCGCGCCTAGCTGTTGTAATGATTTAAT |

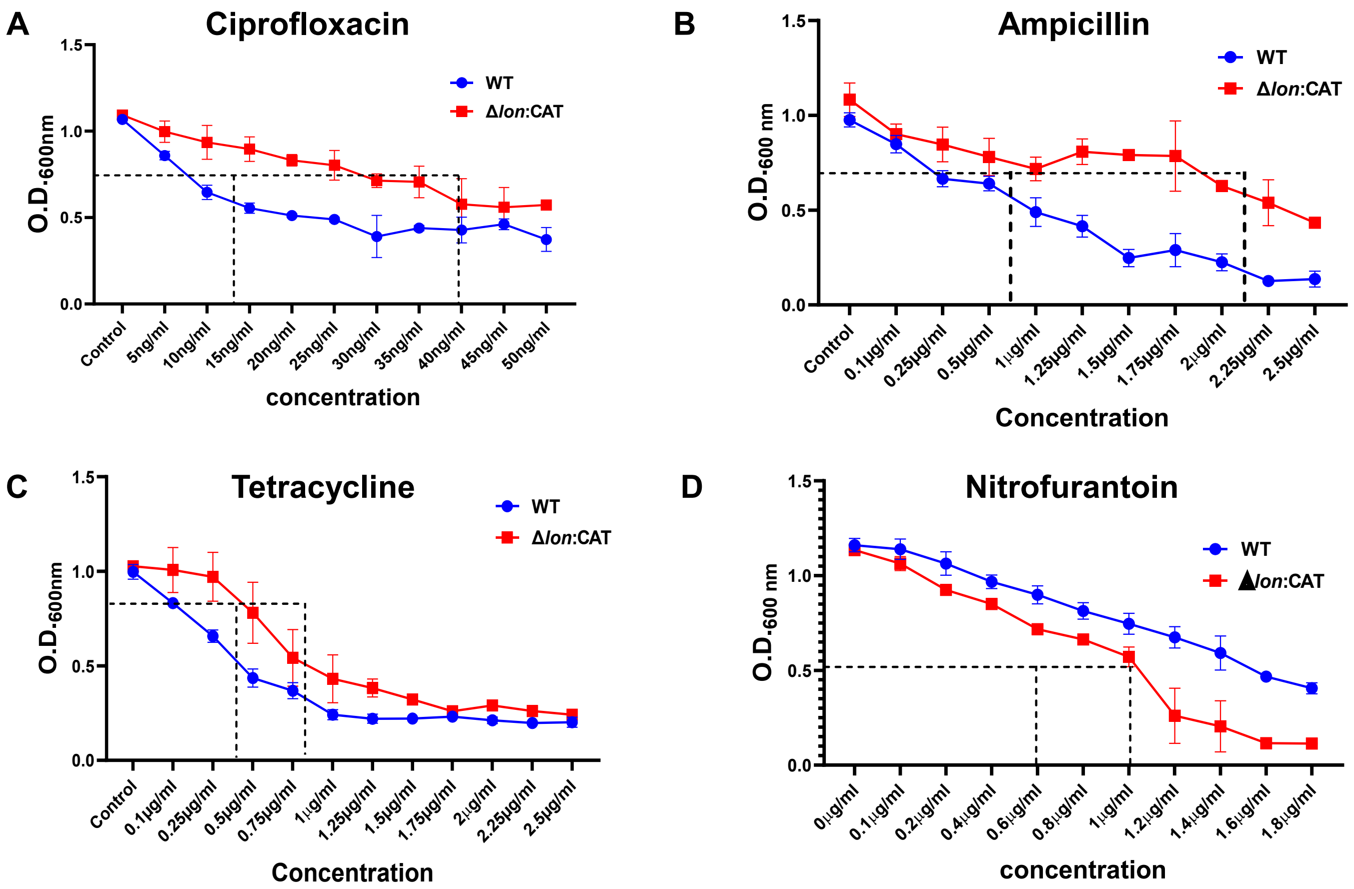

**SI Fig 1: The Δ*lon* strain is resistant to most antibiotics but sensitive to nitrofurantoin.** *E. coli* MG1655 WT and Δ*lon* strains were cultured for 6 hr at 37 °C and 160 rpm in the presence of different antibiotics. MIC broth dilution assay with (A) ciprofloxacin , (B) Ampicillin, (C) Tetracycline and (D) Nitrofurantoin was performed, and data is represented as line plot. The data are representative of at least three independent experiments plotted as mean ± SD.

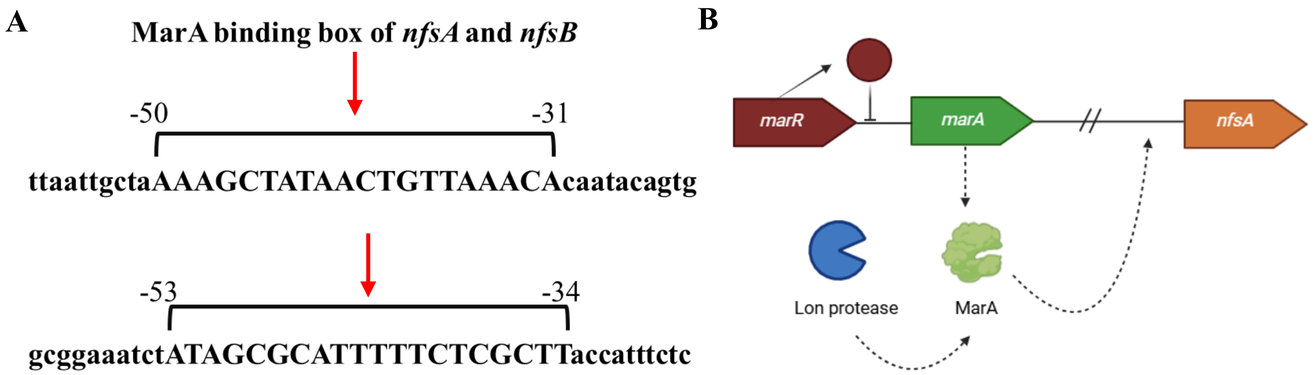

**SI Fig 2. Representative image of Mar A binding box of *nfsA* and *nfsB* :** (A)Position of Site Center Relative to Transcription Start Site for *nfsA* and *nfsB* (bp) are -40.5 and -43.5 respectively (marked with red arrow) .(B) Graphical representation showing role of Lon and MarA in modulating the expression of *nfsA*

A

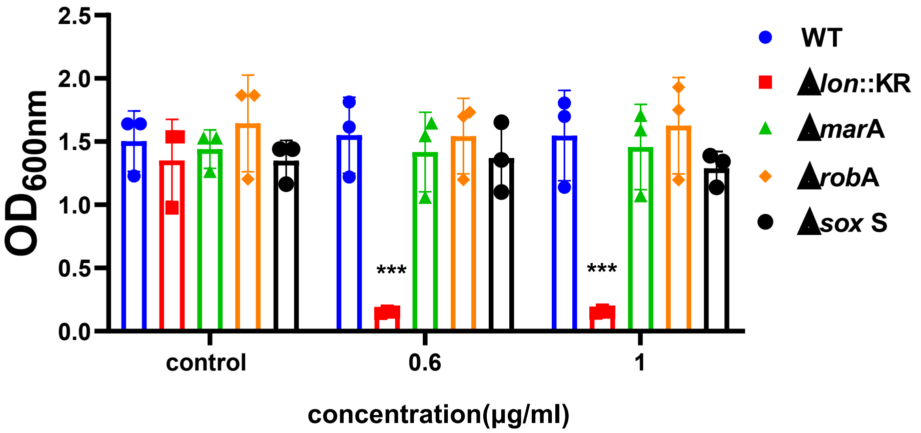

**SI Fig 3. Δ*marA*, Δ*rob* and Δ*soxS* don't show growth reduction upon treatment with nitrofurantoin.** *E. coli* WT, Δ*lon, ΔmarA, Δrob and ΔsoxS* were cultured in the presence of nitrofurantoin at 370 °C and 160 rpm. (A) Growth was assayed by measuring the O.D. at 600 nm using a UV-visible spectrophotometer after growing the cells at different doses of nitrofurantoin for 6 hr. The data are representative of three independent experiments plotted as mean ± SD. Statistical analysis was performed using two-way ANOVA and one-way ANOVA for (D), where * indicates $p<0.05$.

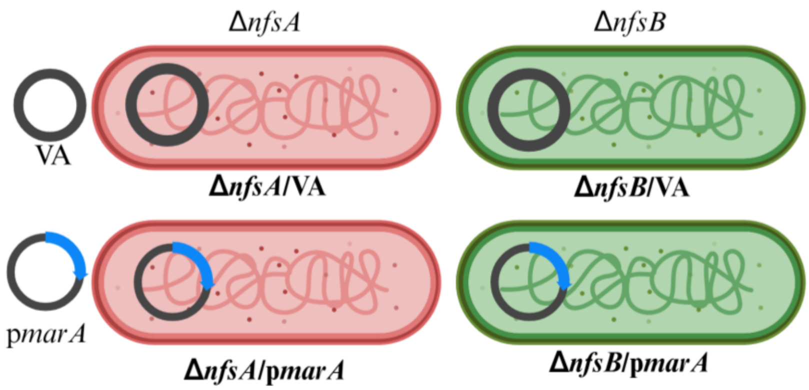

**SI Fig 4. Schematic representation of 4 strains generated through the transformation of p*marA* in Δ*nfsA* and Δ*nfsB***

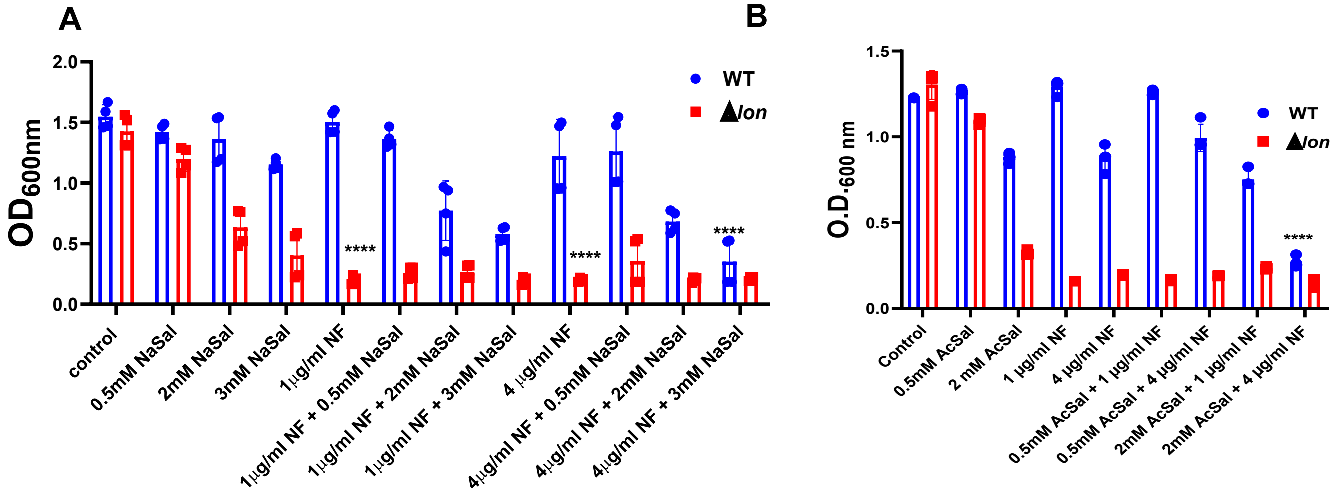

**SI Fig 5 . Combination of NaSal and AcSal with nitrofurantoin induces growth reduction in the WT strain.** *E. coli* MG1655 WT and Δ*lon* were cultured in the presence of nitrofurantoin at 370 °C and 160 rpm. (A) Growth was assayed by measuring the O.D. at 600 nm using a UV-visible spectrophotometer after growing the cells at different doses of nitrofurantoin and NaSal for 6 hr ;(B) Growth was assayed by measuring the O.D. at 600 nm using a UV-visible spectrophotometer after growing the cells at different doses of nitrofurantoin and AcSal for 6 hr; The data represent three independent experiments, plotted as the mean ± SD. Statistical analysis was performed using two-way ANOVA and one-way ANOVA for (D), where * indicates $p<0.05$.

SRI SATHYA SAI INST OF HIGHER MEDICAL SC

bioMerieux Customer:
System #: 8799

### Laboratory Report

Printed Oct 9, 2018 11:55 IST
Printed by: sairam

Patient Name: NI1
Isolate Group: PSNNI1-1

Patient ID: ATCC35218
Bench: URINE

Selected Organism: Escherichia coli

| |
|---|
| **Comments:** |

| **Identification Information** | | | | |
|---|---|---|---|---|
| **Selected Organism** | Escherichia coli | | | |
| | **Entered:** | Oct 7, 2018 11:06 IST | **By:** | sairam |
| **Analysis Messages:** | | | | |

| Susceptibility Information | **Card:** | AST-N280 | **Lot Number:** | 7000524103 | **Expires:** | May 4, 2019 12:00 IST |
|---|---|---|---|---|---|---|
| | **Completed:** | Oct 7, 2018 19:10 IST | **Status:** | Final | **Analysis Time:** | 8.25 hours |

| Antimicrobial | MIC | Interpretation | Antimicrobial | MIC | Interpretation |
|---|---|---|---|---|---|
| Ampicillin | 8 | S | Meropenem | <= 0.25 | S |
| Amoxicillin/Clavulanic Acid | 8 | S | Amikacin | <= 2 | S |
| Piperacillin/Tazobactam | <= 4 | S | Gentamicin | <= 1 | S |
| Cefuroxime | 4 | S | Nalidixic Acid | 4 | S |
| Cefuroxime Axetil | 4 | S | Ciprofloxacin | <= 0.25 | S |
| Ceftriaxone | <= 1 | S | Tigecycline | <= 0.5 | S |
| Cefoperazone/Sulbactam | <= 8 | S | Nitrofurantoin | <= 16 | S |
| Cefepime | <= 1 | S | Colistin | <= 0.5 | S |
| Ertapenem | <= 0.5 | S | Trimethoprim/Sulfamethoxazole | <= 20 | S |
| Imipenem | 0.5 | S | | | |

+= Deduced drug *= AES modified **= User modified

| AES Findings: | **Last Modified:** May 28, 2015 12:17 IST | **Parameter Set:** Copy of CLSI+Natural Resistance |
|---|---|---|
| **Confidence Level:** | Consistent | |

Installed VITEK 2 Systems Version: 06.01
MIC Interpretation Guideline: Copy of CLSI M100-S24 (2014)
AES Parameter Set Name: Copy of CLSI+Natural Resistance

Therapeutic Interpretation Guideline: NATURAL RESISTANCE
AES Parameter Last Modified: May 28, 2015 12:17 IST

SRI SATHYA SAI INST OF HIGHER MEDICAL SC

### Laboratory Report

bioMerieux Customer:
System #: 8799

Printed Apr 11, 2018 11:20 IST
Printed by: sairam

Patient Name: S13
Isolate Group: NS13-1

Patient ID: GE00071982
Bench: URINE

Selected Organism: Escherichia coli

| **Comments:** |
|---|

| **Identification Information** | | | |
|---|---|---|---|
| **Selected Organism** | Escherichia coli | | |
| | **Entered:** Apr 8, 2018 12:18 IST | **By:** | sairam |
| **Analysis Messages:** | | | |

| Susceptibility Information | **Card:** | AST-N280 | **Lot Number:** | 7000524103 | **Expires:** | May 4, 2019 12:00 IST |
|---|---|---|---|---|---|---|
| | **Completed:** | Apr 8, 2018 20:32 IST | **Status:** | Final | **Analysis Time:** | 8.25 hours |

| **Antimicrobial** | **MIC** | **Interpretation** | **Antimicrobial** | **MIC** | **Interpretation** |
|---|---|---|---|---|---|
| Ampicillin | 8 | S | Meropenem | <= 0.25 | S |
| Amoxicillin/Clavulanic Acid | <= 2 | S | Amikacin | 4 | S |
| Piperacillin/Tazobactam | <= 4 | S | Gentamicin | <= 1 | S |
| Cefuroxime | 8 | S | Nalidixic Acid | >= 32 | R |
| Cefuroxime Axetil | 8 | I | Ciprofloxacin | 1 | S |
| Ceftriaxone | <= 1 | S | Tigecycline | <= 0.5 | S |
| Cefoperazone/Sulbactam | <= 8 | S | Nitrofurantoin | <= 16 | S |
| Cefepime | <= 1 | S | Colistin | <= 0.5 | S |
| Ertapenem | <= 0.5 | S | Trimethoprim/Sulfamethoxazole | <= 20 | S |
| Imipenem | <= 0.25 | S | | | |

+= Deduced drug *= AES modified **= User modified

| AES Findings: | **Last Modified:** May 28, 2015 12:17 IST | **Parameter Set:** | Copy of CLSI+Natural Resistance |
|---|---|---|---|
| **Confidence Level:** | Consistent | | |

Installed VITEK 2 Systems Version: 06.01
MIC Interpretation Guideline: Copy of CLSI M100-S24 (2014)
Therapeutic Interpretation Guideline: NATURAL RESISTANCE
AES Parameter Set Name: Copy of CLSI+Natural Resistance
AES Parameter Last Modified: May 28, 2015 12:17 IST

SRI SATHYA SAI INST OF HIGHER MEDICAL SC

bioMerieux Customer: Laboratory Report Printed Oct 10, 2018 10:09 IST
System #: 8799 Printed by: sairam

Patient Name: EC1 Patient ID: PN00048158
Isolate Group: PSNEC1-1 Bench: URINE

Selected Organism: Escherichia coli

| |
|---|
| **Comments:** |

| **Identification Information** | | | | |
|---|---|---|---|---|
| **Selected Organism** | Escherichia coli | | | |
| | **Entered:** | Oct 9, 2018 12:16 IST | **By:** | sairam |
| **Analysis Messages:** | | | | |

| Susceptibility Information | **Card:** | AST-N280 | **Lot Number:** | 7000524103 | **Expires:** | May 4, 2019 12:00 IST |
|---|---|---|---|---|---|---|
| | **Completed:** | Oct 9, 2018 22:43 IST | **Status:** | Final | **Analysis Time:** | 11.25 hours |

| Antimicrobial | MIC | Interpretation | Antimicrobial | MIC | Interpretation |
|---|---|---|---|---|---|
| Ampicillin | >= 32 | R | Meropenem | 8 | *R |
| Amoxicillin/Clavulanic Acid | >= 32 | R | Amikacin | >= 64 | R |
| Piperacillin/Tazobactam | >= 128 | R | Gentamicin | >= 16 | R |
| Cefuroxime | >= 64 | R | Nalidixic Acid | >= 32 | R |
| Cefuroxime Axetil | >= 64 | R | Ciprofloxacin | >= 4 | R |
| Ceftriaxone | >= 64 | R | Tigecycline | <= 0.5 | S |
| Cefoperazone/Sulbactam | >= 64 | R | Nitrofurantoin | >= 512 | R |
| Cefepime | >= 64 | R | Colistin | 2 | S |
| Ertapenem | >= 8 | R | Trimethoprim/Sulfamethoxazole | >= 320 | R |
| Imipenem | >= 16 | R | | | |

+= Deduced drug *= AES modified **= User modified

| AES Findings: | | **Last Modified:** | May 28, 2015 12:17 IST | **Parameter Set:** | Copy of CLSI+Natural Resistance |
|---|---|---|---|---|---|
| **Confidence Level:** | Consistent | | | | |
| **Phenotype:** | BETA-LACTAMS | ESBL + CARBAPENEMASE (METALLO- OR KPC),RESISTANT CARBAPENEMS (IMPERMEABILITY) | | | |
| | AMINOGLYCOSIDES | RESISTANT GEN TOB NET AMI (AAC(6')+?) | | | |

Installed VITEK 2 Systems Version: 06.01
MIC Interpretation Guideline: Copy of CLSI M100-S24 (2014) Therapeutic Interpretation Guideline: NATURAL RESISTANCE
AES Parameter Set Name: Copy of CLSI+Natural Resistance AES Parameter Last Modified: May 28, 2015 12:17 IST

SRI SATHYA SAI INST OF HIGHER MEDICAL SC

bioMerieux Customer:
System #: 8799

### Laboratory Report

Printed Oct 10, 2018 10:10 IST
Printed by: sairam

Patient Name: EC3
Isolate Group: PSNEC3-1

Patient ID: PN00289627
Bench: URINE

Selected Organism: Escherichia coli

| |
|---|
| **Comments:** |

| | | | | |
|---|---|---|---|---|
| **Identification Information** | | | | |
| **Selected Organism** | Escherichia coli | | | |
| | **Entered:** | Oct 9, 2018 12:16 IST | **By:** | sairam |
| **Analysis Messages:** | | | | |

| Susceptibility Information | | | | | |
|---|---|---|---|---|---|
| | **Card:** AST-N280 | | **Lot Number:** 7000524103 | **Expires:** | May 4, 2019 12:00 IST |
| | **Completed:** Oct 9, 2018 17:42 IST | | **Status:** Final | **Analysis Time:** | 6.25 hours |
| **Antimicrobial** | **MIC** | **Interpretation** | **Antimicrobial** | **MIC** | **Interpretation** |
| Ampicillin | >= 32 | R | Meropenem | >= 16 | R |
| Amoxicillin/Clavulanic Acid | >= 32 | R | Amikacin | 16 | S |
| Piperacillin/Tazobactam | >= 128 | R | Gentamicin | 4 | S |
| Cefuroxime | >= 64 | R | Nalidixic Acid | >= 32 | R |
| Cefuroxime Axetil | >= 64 | R | Ciprofloxacin | >= 4 | R |
| Ceftriaxone | >= 64 | R | Tigecycline | <= 0.5 | S |
| Cefoperazone/Sulbactam | >= 64 | R | Nitrofurantoin | >= 512 | R |
| Cefepime | >= 64 | R | Colistin | 2 | S |
| Ertapenem | >= 8 | R | Trimethoprim/Sulfamethoxazole | >= 320 | R |
| Imipenem | >= 16 | R | | | |

+= Deduced drug *= AES modified **= User modified

| AES Findings: | | **Last Modified:** May 28, 2015 12:17 IST | **Parameter Set:** Copy of CLSI+Natural Resistance |
|---|---|---|---|
| **Confidence Level:** | Consistent | | |
| **Phenotype:** | BETA-LACTAMS | RESISTANT CARBAPENEMS (IMPERMEABILITY),ESBL + CARBAPENEMASE (METALLO- OR KPC) | |
| | AMINOGLYCOSIDES | RESISTANT TOB NET AMI (AAC(6')) | |

Installed VITEK 2 Systems Version: 06.01
MIC Interpretation Guideline: Copy of CLSI M100-S24 (2014)
Therapeutic Interpretation Guideline: NATURAL RESISTANCE
AES Parameter Set Name: Copy of CLSI+Natural Resistance
AES Parameter Last Modified: May 28, 2015 12:17 IST
